# PANOPLY: A cloud-based platform for automated and reproducible proteogenomic data analysis

**DOI:** 10.1101/2020.12.04.410977

**Authors:** D. R. Mani, Myranda Maynard, Ramani Kothadia, Karsten Krug, Karen E. Christianson, David Heiman, Karl R. Clauser, Chet Birger, Gad Getz, Steven A. Carr

## Abstract

Proteogenomics involves the integrative analysis of genomic, transcriptomic, proteomic and post-translational modification data produced by next-generation sequencing and mass spectrometry-based proteomics. Several publications by the Clinical Proteomic Tumor Analysis Consortium (CPTAC) and others have highlighted the impact of proteogenomics in enabling deeper insight into the biology of cancer and identification of potential drug targets. In order to encapsulate the complex data processing required for proteogenomics, and provide a simple interface to deploy a range of algorithms developed for data analysis, we have developed PANOPLY—a cloud-based **p**latform for **a**utomated a**n**d repr**o**ducible **p**roteogenomic data ana**ly**sis. A wide array of algorithms have been implemented, and we highlight the application of PANOPLY to the analysis of cancer proteogenomic data.

## PANOPLY

Proteogenomics involves the integrative analysis of genomic, transcriptomic, proteomic and post-translational modification (PTM) data produced by next-generation sequencing and mass spectrometry-based proteomics. Several publications by the Clinical Proteomic Tumor Analysis Consortium (CPTAC) and others have highlighted the impact of proteogenomics in enabling deeper insight into the biology of cancer and identification of potential drug targets^1–4^. In order to encapsulate the complex data processing required for proteogenomics, and provide a simple interface to deploy a range of algorithms developed for data analysis, we have developed PANOPLY—a cloud-based **p**latform for **a**utomated a**n**d repr**o**ducible **p**roteogenomic data ana**ly**sis. A wide array of algorithms have been implemented, and we highlight the application of PANOPLY to the analysis of cancer proteogenomic data..

PANOPLY uses state-of-the-art statistical and machine learning algorithms to transform multi-omic data from cancer samples into biologically meaningful and interpretable results. PANOPLY provides a comprehensive collection of proteogenomic data analysis methods including sample QC (sample quality evaluation using profile plots and tumor purity scores^1^, identify sample swaps, etc.), association analysis, RNA and copy number correlation (to proteome), connectivity map (CMAP) analysis^5^, outlier analysis using BlackSheep^6^, PTM Signature Enrichment Analysis (PTM-SEA)^7^, Gene Set Enrichment Analysis (GSEA)^8^ and single-sample GSEA^7^, consensus clustering, and multi-omic clustering using non-negative matrix factorization (NMF) (**Figure 1A**). Most analysis modules include a report generation task that outputs an interactive HTML report summarizing results from the analysis (**Figure 1B**). A complete list of PANOPLY task modules with documentation can be found at the PANOPLY wiki (https://github.com/broadinstitute/PANOPLY/wiki). The types of liquid chromatography-tandem mass spectrometry-based (LC-MS/MS) proteomics data that are amenable to analysis by PANOPLY includes isobaric mass tag label-based LC-MS/MS approaches like iTRAQ, TMT and TMTPro profiling of the proteome and multiple PTM-omes including phospho-, acetyl- and ubiquitylomes.

**Figure 1.**
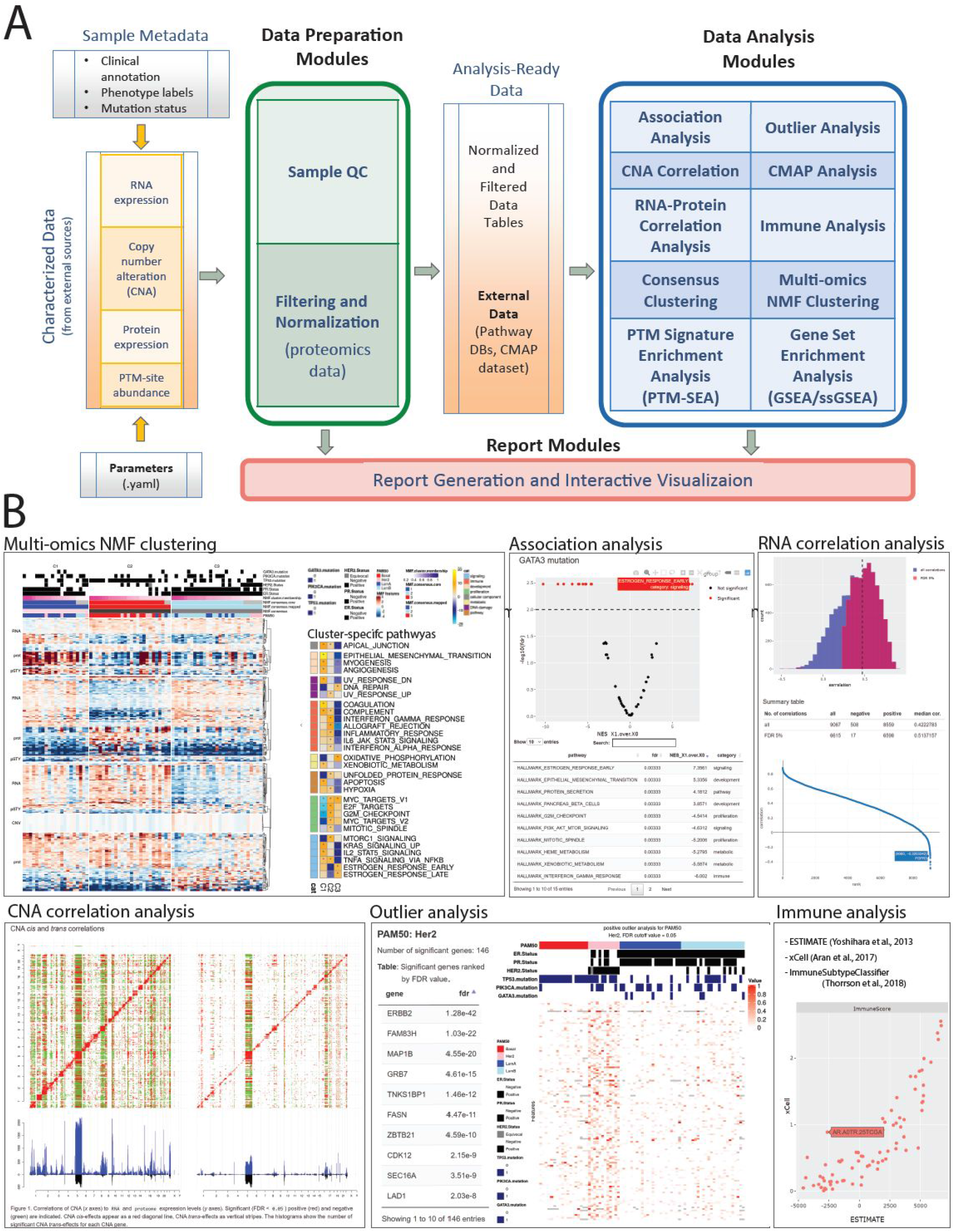
Overview of PANOPLY architecture and the various tasks that constitute the complete workflow. Tasks can also be run independently, or combined into custom workflows, including new tasks added by users. (**A**) Inputs to PANOPLY consists of (i) externally characterized genomics and proteomics data (in gct format); (ii) sample phenotypes and annotations (in csv format); and (iii) parameters settings (in yaml format). Panoply modules are grouped into *Data Preparation Modules* (green box), *Data Analysis Modules* (blue box) and *Report Modules* (red box). Data preparation modules perform quality checks on input data followed by optional normalization and filtering for proteomics data. Analysis ready data tables are then used as inputs to the data analysis modules. Results from the data analysis modules are summarized in interactive reports generated by appropriate report modules. (**B**) A sampling of reports from PANOPLY for the tutorial BRCA dataset ^1^ that can be explored in a web browser. The full suite of reports (html) derived from the tutorial dataset can be accessed and explored at http://prot-shiny-vm.broadinstitute.org:3838/panoply-tutorial/reports, with an interpretation summary at https://github.com/broadinstitute/PANOPLY/wiki/Navigating-Results. <u>Multi-omics NMF clustering:</u> Heatmap (left) depicts multi-omics expression patterns across three NMF clusters derived from integrative clustering of copy-number alterations (CNA), RNA, protein and phosphosite data. Annotation tracks depict mutation status of key BRCA genes (*TP53, GATA3, PIK3CA*), hormone receptor status (ER, PR, HER2) and PAM50 transcriptional subtypes. Right heatmap shows activity scores of cancer hallmark pathways for the three NMF clusters. <u>Association analysis:</u> Interactive volcano plot illustrating differentially abundant signatures of cancer hallmark pathways between tumors with *GATA3* mutations (right side) compared to *GATA3* wild-type tumors (left side). A browseable table summarizes the results. <u>RNA correlation analysis:</u> Histogram of all (blue) and significant (red, FDR < 0.05) gene-level Pearson correlations between RNA and protein data. A table summarizing high-level results and an interactive correlation (y-axis) rank (x-axis) plot enables further exploration of the results. <u>CNA correlation analysis:</u> pairwise gene-level correlations between CNA (x-axis in both panels) and RNA (y-axis, left panel) and proteome (y-axis, right panel). Significant (FDR < 0.05) positive (red) and negative (green) correlations are indicated. CNA *cis*-effects appear as a red diagonal line, CNA *trans*-effects as vertical stripes. The histograms show the number of significant CNA *trans*-effects for each CNA gene. <u>Outlier analysis:</u> Phospho-level outlier analysis showing outliers in the HER2 PAM50 subtype in a searchable table and accompanying heatmap. <u>Immune analysis:</u> Computational approaches to infer immune/stromal tumor components, and immune subtype from RNA expression data are implemented in PANOPLY. Shown are immune scores derived from ESTIMATE^9^ (x-axis) and xCell^10^ (y-axis) for 77 BRCA tumors.

PANOPLY leverages Terra (http://app.terra.bio)—a Google Cloud-based cloud-native platform for extreme-scale data analysis, sharing, and collaboration—to host proteogenomic workflows, and is designed to be flexible, automated, reproducible, scalable, and secure. Using the underlying architecture of Terra, PANOPLY analysis algorithms and methods are implemented as *tasks*, interchangeably referred to as *modules*. A series of tasks constitutes a *workflow*, which represents a pipeline with outputs of one or more tasks feeding into inputs of downstream tasks. Tasks and workflows are defined using the Workflow Description Language (WDL), allowing users to customize pipelines and quickly add their own tasks to the PANOPLY workflow. The central element of PANOPLY is the Terra *workspace*—a shareable collection of everything needed for a data analysis project, including data located on cloud storage linked with the workspace, workflows encapsulating algorithms and pipelines, analysis parameters, results, and a data model that organizes sample meta-data into collections of participants, samples, and sample sets. With versioned workflows, workspaces enable reproducible research by completely encapsulating input data, analysis methods, settings, and results in a shareable cloud-based space.

PANOPLY v1.0 consists of the following components:

- A workspace (https://app.terra.bio/#workspaces/broad-firecloud-cptac/PANOPLY_Production_Pipelines_v1_0) with a preconfigured unified workflow to automatically run all analysis tasks on proteomics (global proteome, phosphoproteome, acetylome, ubiquitylome), transcriptome and copy number data; and an additional workspace (https://app.terra.bio/#workspaces/broad-firecloud-cptac/PANOPLY_Production_Modules_v1_0) that includes separate methods for each analysis component.
- An interactive Jupyter notebook that provides step-by-step instructions for uploading data, identifying data types, specifying parameters, and setting up the PANOPLY workspace for analyzing a new dataset.
- A GitHub repository (https://github.com/broadinstitute/PANOPLY) with code, documentation and description of algorithms.
- A tutorial (https://github.com/broadinstitute/PANOPLY/wiki/PANOPLY-Tutorial) illustrating the application of PANOPLY (https://app.terra.bio/#workspaces/broad-firecloud-cptac/PANOPLY_Tutorial) to a published breast cancer dataset and demonstrating the practical relevance of PANOPLY by regenerating many of the results described in (Mertins et al)^1^ with minimal effort.

PANOPLY has been used for analyzing data from many CPTAC and other cancer proteogenomic landscape studies, and workspaces with results for published data sets, including CPTAC BRCA^1,4^ (https://app.terra.bio/#workspaces/broad-firecloud-cptac/PANOPLY_CPTAC_BRCA) and LUAD^3^ (https://app.terra.bio/#workspaces/broad-firecloud-cptac/PANOPLY_CPTAC_LUAD) are available. While PANOPLY has been developed and presented in the context of cancer proteogenomic studies, it can be applied to any proteogenomics study with appropriate data, vastly simplifying the analysis and integration process. While the current release of PANOPLY contains algorithms for integrative proteogenomic data analysis, future versions will include workflows to perform raw data characterization of genomics and proteomics data. New analysis methods from recent CPTAC proteogenomic studies will also be added in new releases. We envision that use of PANOPLY will provide a quick and comprehensive baseline analysis for proteogenomic studies, leading to many disease specific hypotheses that can be explored further using additional computational and wet-lab experiments.

## Acknowledgements

This work was supported by grants from the National Cancer Institute (NCI) Clinical Proteomic Tumor Analysis Consortium grants NIH/NCI U24CA210979 (to DRM and GG) and NIH/NCI U24-CA210986 and NIH/NCI U01 CA214125 (to SAC).

## References

1. Mertins, P. et al. Proteogenomics connects somatic mutations to signalling in breast cancer. Nature 534, 55–62 (2016).

2. Zhang, B. et al. Proteogenomic characterization of human colon and rectal cancer. Nature 513, 382–387 (2014).

3. Gillette, M. A. et al. Proteogenomic Characterization Reveals Therapeutic Vulnerabilities in Lung Adenocarcinoma. Cell 182, 200–225.e35 (2020).

4. Krug, K. et al. Proteogenomic Landscape of Breast Cancer Tumorigenesis and Targeted Therapy. Cell 183, 1–21 (2020).

5. Subramanian, A. et al. A Next Generation Connectivity Map: L1000 Platform and the First 1,000,000 Profiles. Cell 171, 1437–1452.e17 (2017).

6. Blumenberg, L. et al. BlackSheep: A Bioconductor and Bioconda package for differential extreme value analysis. doi:10.1101/825067.

7. Krug, K. et al. A Curated Resource for Phosphosite-specific Signature Analysis. Mol. Cell. Proteomics 18, 576–593 (2019).

8. Subramanian, A. et al. Gene set enrichment analysis: a knowledge-based approach for interpreting genome-wide expression profiles. Proc. Natl. Acad. Sci. U. S. A. 102, 15545–15550 (2005).

9. Yoshihara, K. et al. Inferring tumour purity and stromal and immune cell admixture from expression data. Nat. Commun. 4, 1–11 (2013).

10. Aran, D., Hu, Z. & Butte, A. J. xCell: digitally portraying the tissue cellular heterogeneity landscape. Genome Biol. 18, 220 (2017).

