## Supplement for "PANOPLY: A cloud-based platform for automated and reproducible proteogenomic data analysis"

All supplementary information is available online at **GitHub** (https://github.com/broadinstitute/PANOPLY) and **Terra** (https://app.terra.bio/), and these resource URLs are included in the main text. Here we tabulate these for convenient access.

| Category | Item | Location (URL) | Notes |
|---|---|---|---|
| **PANOPLY Release v1.0** | PANOPLY pipelines | https://app.terra.bio/#workspaces/broad-firecloud-cptac/PANOPLY_Production_Pipelines_v1_0 | Terra workspace with pre-configured pipelines, including startup notebook for easy data input and workspace configuration |
| | PANOPLY modules | https://app.terra.bio/#workspaces/broad-firecloud-cptac/PANOPLY_Production_Modules_v1_0 | Terra workspace with all individual modules. This enables users to pick and choose modules and customize execution, but requires more knowledge of setting up inputs. |
| **Tutorial** | Tutorial description | https://github.com/broadinstitute/PANOPLY/wiki/PANOPLY-Tutorial | Step-by-step instruction on running the PANOPLY tutorial |
| | Tutorial workspace | https://app.terra.bio/#workspaces/broad-firecloud-cptac/PANOPLY_Tutorial | Terra workspace with tutorial instructions, data, analysis and results |
| **Case Studies** | BRCA | https://app.terra.bio/#workspaces/broad-firecloud-cptac/PANOPLY_CPTAC_BRCA | Terra workspace with data, analysis and results from the (Krug, et al,., 2020, *Cell*) breast cancer study |
| | LUAD | https://app.terra.bio/#workspaces/broad-firecloud-cptac/PANOPLY_CPTAC_LUAD | Terra workspace with data, analysis and results from the (Gillette, et al,., 2020, *Cell*) lung adenocarcinoma study |
| **Code** | Code for v1.0 release | https://github.com/broadinstitute/PANOPLY/tree/release-1_0 | Code for current release of PANOPLY |
| | Development code repository | https://github.com/broadinstitute/PANOPLY | Includes release and development code |

| | | | |
|---|---|---|---|
| **Methods** | Computational methods used in PANOPLY analysis modules | https://github.com/broadinstitute/PANOPLY/wiki/Data-Analysis-Modules | Documentation for each data analysis module includes a description of methods, with appropriate references |
| | Data pre-processing methods | https://github.com/broadinstitute/PANOPLY/wiki/Data-Preparation-Modules | Documentation for each module includes a description of methods, with appropriate references |
| **Documentation** | PANOPLY documentation | https://github.com/broadinstitute/PANOPLY/wiki | Detailed documentation for each PANOPLY module, including description of method, inputs and outputs |
| | Interpreting PANOPLY reports and results | https://github.com/broadinstitute/PANOPLY/wiki/Navigating-Results | Description of interactive reports generated by PANOPLY in the context of the tutorial dataset |
| | Advanced uses | https://github.com/broadinstitute/PANOPLY/wiki/Customizing-PANOPLY | Customizing PANOPLY and adding new analysis modules |
| | PANOPLY without Terra | https://github.com/broadinstitute/PANOPLY/wiki/PANOPLY-without-Terra | Using PANOPLY on a local computer, without accessing Terra on the cloud. |
